# SAM-DNMT3A, a strategy for induction of genome-wide DNA methylation, identifies DNA methylation as a vulnerability in ER-positive breast cancers

**DOI:** 10.1101/2024.01.16.575955

**Authors:** Mahnaz Hosseinpour, Luis Malaver-Ortega, Laura Perlaza-Jimenez, Jihoon E. Joo, Ling Liu, Elizabeth C. Caldon, Pierre-Antoine Dugué, James G. Dowty, Melissa C. Southey, Joseph Rosenbluh

## Abstract

DNA methylation is an epigenetic mark that plays a critical role in regulation of gene expression. DNA methylase (DNMT) inhibitors, inhibit global DNA methylation, and have been a key tool in studies of DNA methylation in healthy or disease conditions. A major bottleneck is the lack of tools to induce global DNA methylation. Here, we engineered a CRISPR based approach, that was initially designed, to enable site specific DNA methylation. Using the synergistic activation mediator (SAM) system, we unexpectedly found that regardless of the targeted sequence any sgRNA induced global genome-wide DNA methylation. We termed this new method SAM-DNMT3A and show that induction of global DNA methylation is a unique vulnerability in ER-positive breast cancer suggesting a therapeutic approach. Our findings highlight the need of caution when using CRISPR based approaches for inducing DNA methylation and demonstrate a new method for global induction of DNA methylation.

## Introduction

DNA methylation is a reversable epigenetic modification that plays an important role in regulating gene expression and maintaining the stability of the genome^1^. DNA methylation at promoters regulates gene expression by preventing transcription factors from binding to DNA and initiating transcription^2^. In contrast, DNA methylation at gene body regions can have complex effects on gene expression, sometimes enhancing it^3, 4, 5^. Aberrant DNA methylation patterns have been associated with a wide range of diseases, including cancer, neurological disorders, and developmental disorders^1^. Mendelian-like inheritance of DNA methylation has been shown in various organisms^6^. We have shown that in some cases heritable DNA methylation is associated with susceptibility to different cancer types^7, 8, 9^. Although DNA methylation is a fundamental cellular process that is involved in almost every aspect of life, due to the limited available toolkit to study DNA methylation, we still lack comprehensive understanding of how DNA methylation mediates biological processes.

DNA methylation is induced by DNA methylases (DNMT’s) that in humans include *DNMT3A* and *DNMT3B* and is maintained by *DNMT1*^10^. The DNA methylation mark is removed by the *TET1-3* enzymes^11^. Small molecule inhibitors of DNMT’s^12^ (e.g. decitabine and azacytidine) induce global DNA de-methylation and have been used as a powerful tool to study the effects of global DNA de-methylation on normal biological processes such as aging^13^, and disease conditions such as cancer^12^. A major gap in the study of DNA methylation is that we currently lack strategies for induction of global genome-wide DNA methylation.

Clustered Regularly Interspaced Short Palindromic Repeats (CRISPR) technology offers the potential for precise induction of DNA methylation at desired sites in the genome. Using dCas9 fused to *DNMT3A* Liu et al. developed a programable site-specific DNA methylation platform^14^. Subsequent studies found that this system induces low levels of DNA methylation that is prone to off target effects^15^. Pflueger et al.^16^ found that adopting the SunTag system^17^ allows a more specific induction of DNA methylation with less pervasive off-target effects. Here, we show that the SAM system^18^ induces the highest levels of DNA methylation and that induction of DNA methylation with the SAM system induces global genome-wide non-specific DNA methylation. Our results identify SAM-DNMT3A as an effective tool for induction of global DNA methylation and to study the effect of global DNA methylation in normal or disease development.

## Results

### Development of SAM-DNMT3A for induction of DNA methylation

Previous studies using the SunTag system demonstrated that it is more specific than dCas9-DNMT3A fusions for induction of site-specific DNA methylation^16^. Since the SAM system has been successfully utilised for development of highly active CRISPR based programable transcription activators^18^, and since the SAM system requires less plasmids^19^, we developed SAM as an approach for induction of DNA methylation (Fig. 1A). In this approach a lentiviral vector expressing a catalytically inactive Cas9 enzyme (dCas9) fused to a DNMT enzyme, is stably transduced into a cell. Following blasticidine selection for stably integrated cells, a second lentiviral vector expressing a modified sgRNA with two *PP7* RNA binding loops and *DNMT3A*-*PP7* fusion protein, with a puromycin selection gene is introduced (Fig. 1A). This strategy enables recruitment of three DNMTs to a desired site and enables induction of a high degree of DNA methylation (Fig. 1B).

**Figure 1:**
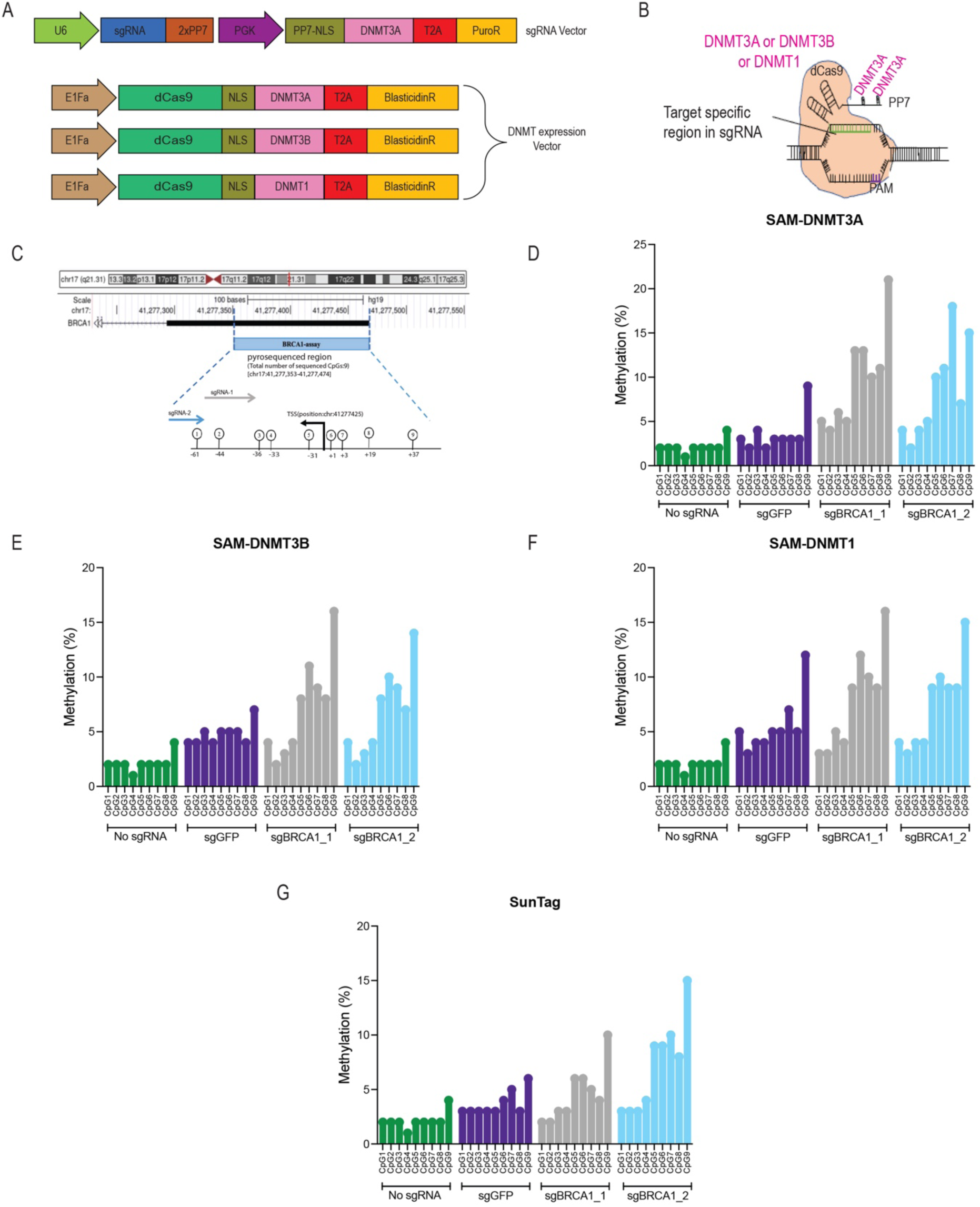
SAM-DNMT3A induces high levels of DNA methylation at desired sites. (A) Vectors used for development of SAM-DNMT3A. The sgRNA vector contains puromycin resistance and a PP7-DNMT3A fusion protein. The DNMT expression vector contains a blasticidine resistance gene and a dCas9 enzyme fused to a DNMT protein. (B) Illustration of SAM-DNMT system. At each target site three DNMT proteins are recruited. (C) sgRNAs used for induction of DNA methylation at the *BRCA1* promoter. (D) DNA methylation measured using Pyrosequencing at CpG sequences of *BRCA1* promoter 3 days following transfection of HEK293T cells with the SAM-DNMT3A system. (E) Like in (D) using SAM-DNMT3B system. (F) Like in (D) using SAM-DNMT1 system. (G) Like in (D) using SunTag-DNMT3A^16^ system.

To test the ability of this approach to induce site specific DNA methylation, we designed two sgRNAs targeting CpG sequences at the *BRCA1* promoter (Fig. 1C). HEK293T cells were transfected with a non-targeting control (sgGFP) or two *BRCA1* targeting sgRNAs fused to either *DNMT3A*, *DNMT3B* or *DNMT1*. Using Pyrosequencing we analysed DNA methylation at the *BRCA1* locus, 3 days post transfection (Fig. 1D-F). Compared to un-transfected cells, even the non-targeting control sgRNAs showed some increase in DNA methylation (Fig. 1D-F). However, for some CpG sites we observed that *BRCA1* targeting sgRNAs induced higher levels of DNA methylation. Of the three DNMT enzymes we found that SAM-DNMT3A induced the highest levels of DNA methylation (Fig. 1D). We further validated SAM-DNTM3A using lentivirus delivered system in K5+/K19-cells^20^, a normal breast immortalised cell line, with two additional genes, *PTEN* and *NF1* (Supplementary Fig. 1A,B). To compare between SunTag and SAM system we used the same *BRCA1* targeting sgRNAs with SunTag system fused to *DNMT3A* (Fig. 1G). We found that the SAM-DNMT3A system induced higher levels of DNA methylation. Based on these experiments we selected the SAM-DNMT3A for all further studies.

### SAM-DNMT3A pooled screens

Pooled genetic screens enable high throughput investigation of genetic perturbations and how they regulate biological processes^21, 22^. We have previously reported that some methylation sites are heritable and some of these heritable methylation sites are associated with increased risk of breast^7, 8^ and prostate^8, 9^ cancer. Increased cell proliferation is a hallmark of cancer^23^ and is associated with many cancer risk associated genes^24^. To identify if any of these heritable methylation marks induce a cancer related phenotype, we designed a pooled sgRNA library that targets the top 1,000 heritable methylation marks and used this library to identify sites in the human genome that upon methylation induce a proliferation phenotype (Supplementary Table S1).

Since SAM-DNMT3A is directed to the gene promoter it is possible that the observed effect will be related to inhibition of transcription (CRISPRi) rather than induction of DNA methylation. To avoid possible CRISPRi effects of SAM-DNMT3A as a control we did the same screen using a catalytically inactive DNMT3A (SAM-DNMT3A-inactive). The High-resolution melting (HRM) assay quantifies the difference in melting curves following bisulfite conversion as a measure of DNA methylation^25^. Using the HRM assay on cells transfected with SAM-DNMT3A or SAM-DNMT3A-inactive with an sgRNA targeting the *BRCA1* promoter, we found that consistent with previous reports^16^, active DNMT3A is required for induction of DNA methylation (Supplementary Fig. 2). Our sgRNA library includes 10,286 sgRNA targeting 1,009 genomic regions in the human genome (Supplementary Table S1). For each of these regions we designed 10 sgRNAs located at the centre of the identified DNA methylation peak (Supplementary Table S1). In addition, we included 737 negative control sgRNAs targeting non-human genes or 262 sgRNAs targeting the *AAVS1* region. MCF7, T47D or BRE80-T5 cells expressing SAM-DNMT3A or SAM-DNMT3A-inactive were transduced with a lentiviral sgRNA library at a multiplicity of infection (MOI) of 0.3, to ensure one sgRNA/cell. Following selection cells were cultured for 21 days and DNA extracted from these cells was used for quantification of sgRNA abundance (Fig. 2A). The effect of an sgRNA on cell proliferation is calculated by comparing sgRNA abundance in SAM-DNMT3A and SAM-DNMT3A-inactive (Supplementary Table S2).

**Figure 2:**
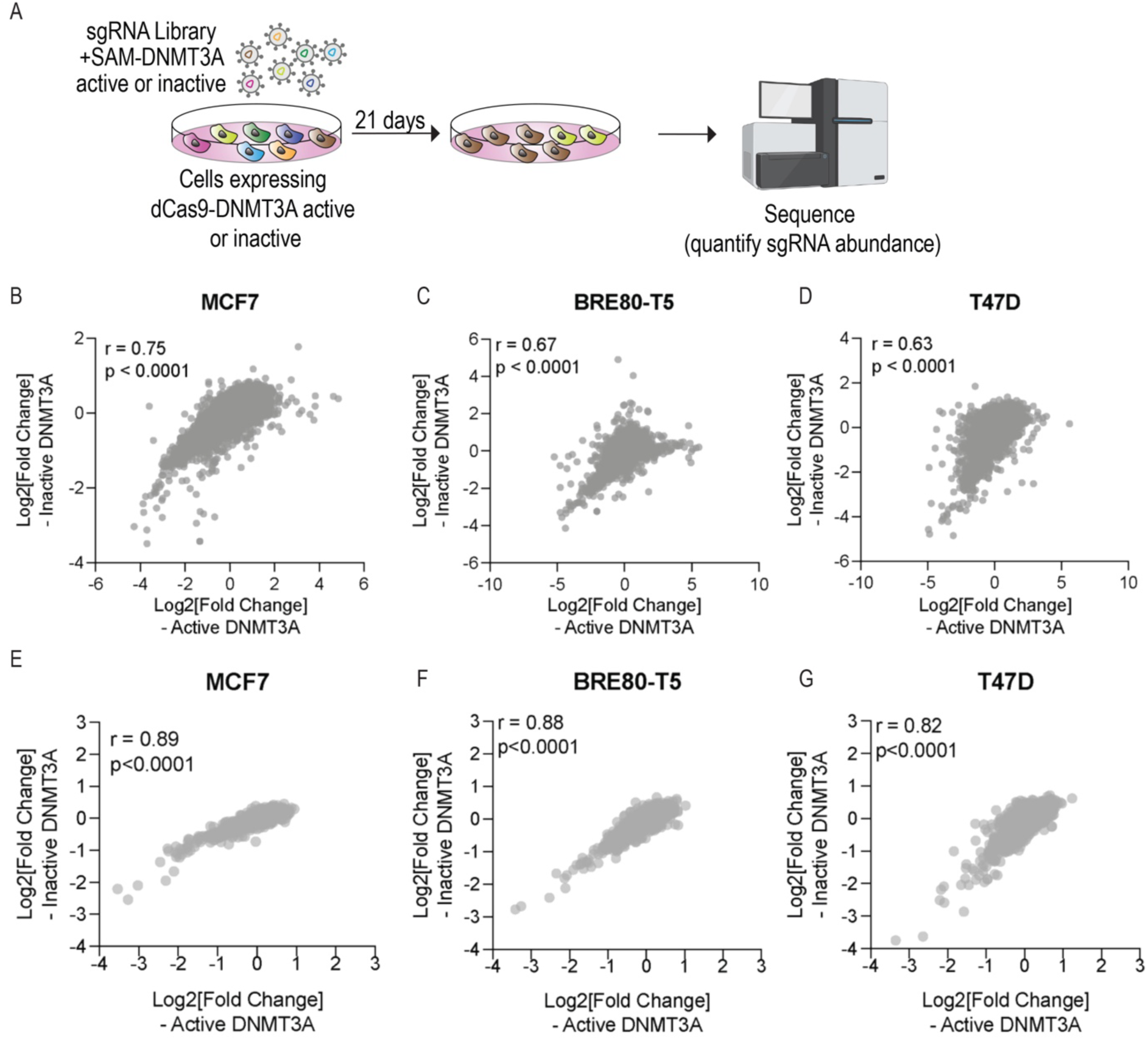
SAM-DNMT3A pooled screens do not identify any hits associated with proliferation. (A) Illustration of the pooled screen strategy used to identify DNA methylation sites that effect cell proliferation. Correlation of sgRNA abundance between cells with active and inactive DNMT3A for (B) MCF7 (C) BRE80-T5 (D) T47D. Correlation of gene scores between cells with active and inactive DNMT3A for (E) MCF7 (F) BRE80-T5 (G) T47D.

We used this approach to screen three breast cell lines. MCF7 and T47D, two ER-positive breast cancer cell lines, and BRE80-T5, a normal immortalised ER-negative mammary epithelial cell line^20^. By comparing sgRNA abundance in cells with SAM-DNMT3A and SAM-DNMT3A-inactive we found no significant difference (Fig. 2B-D and Supplementary Table S2). Using the MAGeCK algorithm^26^ we calculated a gene score for each of these regions and found no significant difference between cells with active or inactive DNMT3A (Fig. 2E-G and Supplementary Table S2). These results indicate that either none of these methylated regions have a functional consequence or that site specific DNA methylation induced with this system is non-specific.

### SAM-DNMT3A induces genome-wide global non-specific DNA methylation

Since we did not find any hits in a pooled proliferation screen we measured the specificity of the SAM-DNMT3A system. Using the EPIC v2.0 array we quantified the levels of DNA methylation in BRE80-T5 or T47D cells containing SAM-DNMT3A or SAM-DNMT3A-inactive. Following transduction with no sgRNA or an sgRNAs targeting *AAVS1*, safe harbour region, or sgRNAs targeting regions in the genome containing hereditable DNA methylation marks^7^. Infected cells were selected with puromycin and 7 days post transduction, genomic DNA extracted from these cells was used for bisulfite conversion and quantification of DNA methylation throughout the genome using the Infinium Methylation EPIC v2.0 microarray (Fig. 3A).

**Figure 3:**
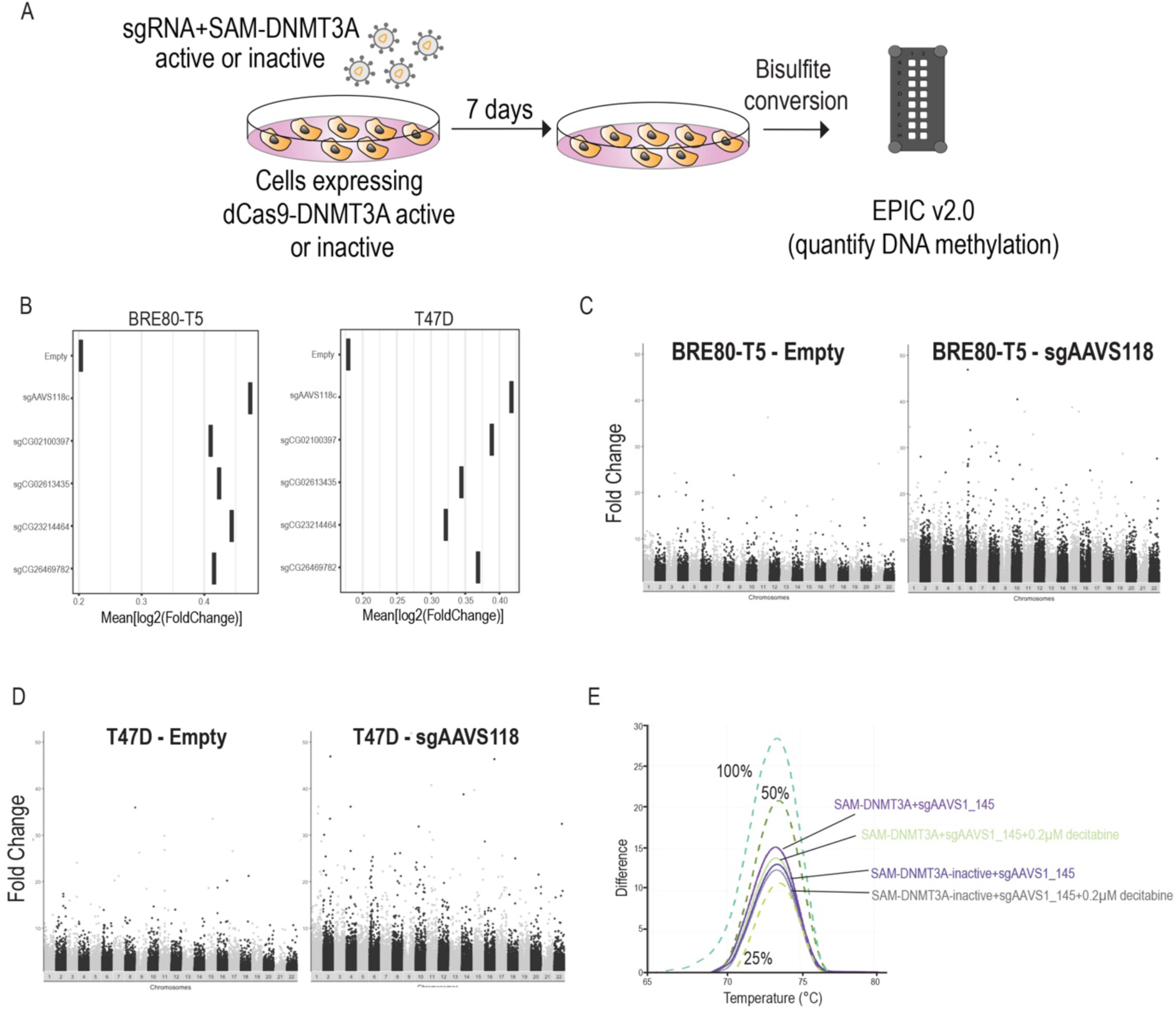
SAM-DNMT3A induces global non-specific DNA methylation. (A) Illustration of experiment aimed to identify global methylation changes induced by SAM-DNMT3A. (B) For each methylation site the fold change in methylation was calculated by comparing methylation levels in cells with SAM-DNMT3A and SAM-DNMT3A-inactive. The average fold change in DNA methylation is plotted for each sgRNA. (C) Manhattan plot showing all methylation sites in BRE80-T5 cells with no sgRNAs or with sgAAVS1_118. (D) Manhattan plot showing all methylation sites in T47D cells with no sgRNAs or with sgAAVS1_118. (E) HRM assay using LINE-1 probes in MCF7 cells expressing SAM-DMT3A or SAM-DNMT3A-inactive with sgAAVS1 in cells treated for 48h with 0.2µM of the DNMT inhibitor decitabine.

For each CpG site in the genome we calculated the fold change in DNA methylation by comparing DNA methylation in cells containing SAM-DNMT3A and cells with SAM-DNMT3A-inactive. Using the mean fold change of global DNA methylation across all sites in the human genome we found that compare to cells with no sgRNA cells expressing a targeting sgRNA directed to any region of the genome had a strong effect on induction of DNA methylation throughout the genome (Fig. 3B). In both BRE80-T5 cells and T47D cells DNA methylation was increased compared to the non-targeting control (Fig. 3B). The increase in DNA methylation was observed throughout the genome and was not restricted to a specific site or chromosome (Fig. 3C,D and Supplementary Fig. 3A,B). To further confirm that the global methylation induced by SAM-DNMT3A is due to the DNMT3A activity we used the DNMT inhibitor decitabine. MCF7 cells expressing SAM-DNMT3A or SAM-DNMT3A-inactive and an sgRNA targeting *AAVS1* were treated with 0.2µM of decitabine and global DNA methylation was monitored using the HRM assay with LINE-1 probes (Fig. 3E). We found that like what we observed using the EPIC array SAM-DNMT3A with an *AAVS1* targeting sgRNA induced an increase in LINE-1 methylation that was supressed by decitabine. Based on these results we conclude that in the presence of an sgRNA the SAM-DNMT3A system induces non-specific global genome DNA methylation that is not related to the sgRNA identity or cell line used.

### Global induction of DNA methylation is a vulnerability in ER-positive breast cancer

During the above-described experiments, we noticed that ER-positive breast cancer cell lines expressing the SAM-DNMT3A system with any sgRNA were difficult to culture for long periods suggesting that induction of global DNA methylation is a vulnerability in ER-positive breast cancers. To test this hypothesis, we used a panel of 7 breast cancer cell lines (4 ER-positive and 3 ER-negative) expressing SAM-DNMT3A or SAM-DNMT3A-inactive with no sgRNA or two *AAVS1* targeting sgRNAs that induce global DNA methylation (Fig. 4A,B). We found that induction of genome wide DNA methylation had no effect on proliferation in ER-negative breast cancers. Conversely, in ER-positive breast cancer cell lines induction of DNA methylation significantly inhibited cell proliferation (Fig. 4C).

**Figure 4:**
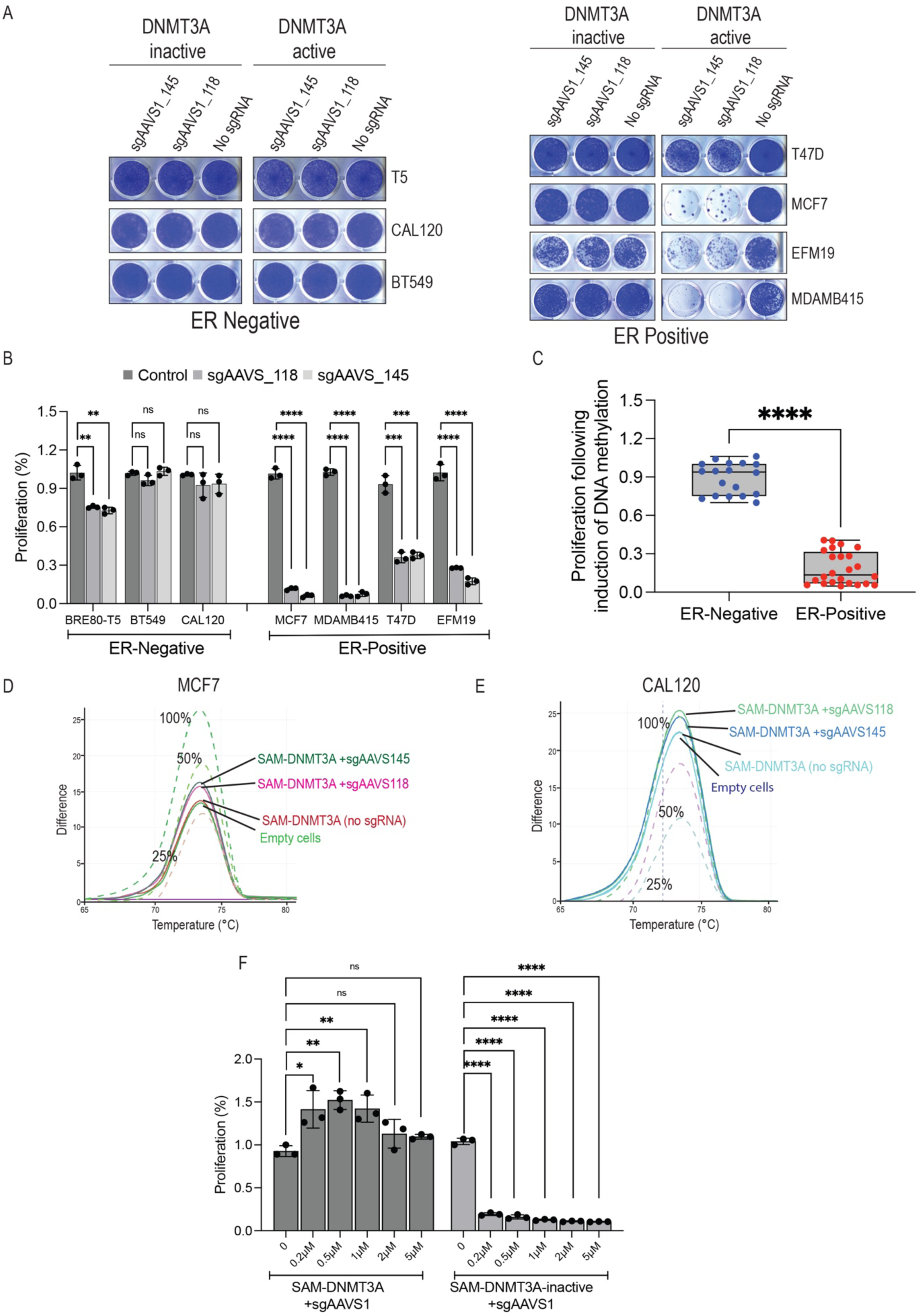
Induction of DNA methylation is a vulnerability in ER positive breast cancers. (A) Images of Crystal violet staining of ER positive and ER negative breast cancer cell lines containing active or inactive DNMT3A 7 days post transduction with no sgRNA or AAVS1 targeting sgRNAs. (B) Quantification of crystal violet images. Unpaired two tailed T.test of three biological replicates was used to calculate a pValue. (C) Proliferation changes of all AAVS1 sgRNAs in ER-positive and ER-negative breast cancers. Unpaired two tailed T.test was used to calculate pValue. (D) Global DNA methylation measured using the HRM assay with probes for LINE elements in MCF7 cells transduced with SAM-DNMT3A and control or AAVS1 targeting sgRNAs. Dotted lines are DNA methylation controls generated using a mix of fully methylated and un-methylated genomic DNA at the indicated ratio. (E) Like (D) in CAL-120 (ER-negative) cell line. (F) Proliferation of MCF7 cells with SAM-DNMT3A or SAM-DNMT3A-inactive and an AAVS1 targeting sgRNA treated with the indicated concentrations of the DNMT inhibitor decitabine. Unpaired two tailed T.test of three replicates was used to calculate a pValue.

To assess DNA methylation induced by SAM-DNMT3A in ER-positive and ER negative breast cancer cells we used the HRM assay with probes measuring DNA methylation at LINE-1 sequences. We found that, as previously reported^27^, the basal levels of DNA methylation were lower in ER-positive breast cancers (Fig. 4D and Supplementary Fig. 4A,B) compared with ER-negative breast cancers (Fig. 4E). In both breast cancer cell types, we observed a ∼10% increase in global DNA methylation induced by SAM-DNMT3A. We further confirmed using the HRM assay that DNA methylation induction was dependent on active DNMT3A (Supplementary Fig. 4A,B). To further validate that reduction in proliferation we observed in ER-positive breast cancer cells is due to global induction of DNA methylation we used a recue experiment. MCF7 (ER-positive) cells expressing SAM-DNMT3A or SAM-DNMT3A-inactive with an *AAVS1* targeting sgRNA were treated with increasing concentrations of the DNMT inhibitor decitabine (Fig. 4F and Supplementary Fig. 4C). High concentrations of decitabine were toxic in all conditions (Supplementary Fig. 4C), however, low concentrations of decitabine rescued growth of MCF7 cells containing the SAM-DNMT3A (Fig. 4F and Supplementary Fig. 4C) demonstrating that SAM-DNMT3A induced proliferation arrest is due to induction of global DNA methylation. These observations demonstrate that SAM-DNMT3A induces global DNA methylation in a cell line independent manner and that induction of DNA methylation is a vulnerability in ER-positive breast cancers.

## Discussion

DNA methylation is the most prevalent epigenetic mark in the human genome. It is a key regulator of normal biological processes and is deregulated in various disease phenotypes^1^. DNA methylation is a fundamental epigenetic modification. It is one of the first epigenetic marks that is induced in the developing stem cell and is critical for lineage commitment^28^. Furthermore, DNA methylation at specific loci could be inherited to off-springs and in some cases is associated with disease risk^7, 9^. Global induction or suppression of DNA methylation is a common cellular mechanism to regulate gene expression and is mediated by DNMT and TET enzymes^1^. Despite the clear importance of understanding how DNA methylation regulates biological processes our current toolbox is limited to DNMT inhibitors. Specifically, current tools to study DNA methylation include strategies for profiling of DNA methylation in cells and tissues and methods to perturb DNA methylation and assess the phenotypic outcomes. The global DNA methylation status in the human genome is measured using bisulfite conversion linked to microarrays or sequencing platforms^29^. For functional studies of DNA methylation DNMT inhibitors are commonly used to induce global DNA de-methylation^12^. A major gap is the lack of tools to induce global DNA methylation. Here, we show that SAM-DNMT3A is a simple robust and effective strategy for induction of global DNA methylation. In the presence of an sgRNA, SAM-DNMT3A induces rapid high-level, genome-wide DNA methylation. SAM-DNMT3A is an important tool for studies of how DNA methylation is associated with phenotypes in normal or disease conditions.

Our results show that SAM-DNMT3A induces genome-wide DNA methylation. Using functional pooled screens we found no consistent phenotypes with SAM-DNMT3A and in global methylation profiling studies we found that any sgRNA we used induced global DNA methylation throughout the genome. Nevertheless, our study finds that CRISPR based systems for induction of DNA methylation are highly prone to off-target effects. Previous studies suggested that systems like SunTag induce higher levels of DNA methylation and are less affected by off-target methylation events^16^. Furthermore, Nunez et al et al.^30^ showed that a transient CRISPR based system induces a highly specific and heritable DNA methylation mark. These reports are in contrast with our findings. It is possible that these approaches observe a more specific effect since they use a transient expression system that is different from our system which uses systemic expression of SAM-DNMT3A. The most likely mechanism of off-target DNA methylation with SAM-DNMT3A is that when the Cas9 sgRNA complex scans the genome to find its target it leaves DNA methylation marks. Since the same mechanism would also apply for other CRISPR based approaches, our results suggest that studies using CRISPR based methylation to silence genes, need to take care and ensure that observed phenotypes are not mediated by an off-target global DNA methylation event.

Using SAM-DNMT3A system we show that ER-positive breast cancers are highly sensitive to induction of DNA methylation. These findings are consistent with previous reports that observed low levels of global DNA methylation in ER-positive breast cancers^27^. Previous studies have suggested that low DNA methylation is important for transcriptional activity of the oestrogen receptor. Here, we show, for the first time, that low levels of DNA methylation are required for proliferation in ER-positive breast cancers and that induction of DNA methylation is a unique vulnerability in these cancers. Although we currently do not have small molecules that induce global DNA methylation these observations suggest that strategies to induce global DNA methylation would be valuable as a treatment in ER-positive breast cancers. These findings could also have implications to ER-positive cancers that are resistant to ER inhibitors. Previous reports suggest that ER-positive breast cancers that are resistant to endocrine therapy have higher levels of DNA methylation^31, 32^. Based on our findings showing ER-positive breast cancers require low levels of DNA methylation suggests that ER dependency and DNA methylation are directly connected and that it is possible that escapers of this process are no longer sensitive to endocrine therapy or to DNA methylation induction. Our work provides a new method for induction of global DNA methylation and suggests that cation should be taken when using CRISPR based approaches to induce site specific DNA methylation.

## Data and reagent availability

All raw and processed data from EPIC arrays are available through GEO (GSE249125). Raw and processed reads from pooled screens are available as Supplementary Tables. All plasmids made in this study are available through Addgene.

## Methods

### Cell lines and media

Cell-line identity was verified by STR at the Australian Genomics Research Facility (AGRF). MCF7 and T47D cells were a gift from Prof. Georgia Chenevix-Trench (QIMR-Berghofer). HEK293FT cells were from Thermo-Fisher (#R70007). All other cell lines used in this study were from ATCC. HEK293FT cell lines were maintained in the complete DMEM medium consisting of Dulbecco’s Modified Eagle Medium (DMEM; Thermo-Fisher Scientific 11965092), 1% Penicillin-Streptomycin (Pen-Strep; Sigma-Aldrich, P4333), 10% Fetal Bovine Serum (FBS; Bovogen, SFBS) and 1% L-glutamin (Sigma-Aldrich, G7513), and supplemented with 1% MEM^™^ Sodium Pyruvate (Thermo-Fisher Scientific, 11360070) and 1% MEM^TM^ Non-Essential^TM^ Amino Acids (Thermo-Fisher Scientific, 11140050). MDAMB415 cell lines were cultured in the complete DMEM medium with addition of 0.01 mg/ml insulin from bovine pancreas (Sigma-Aldrich, 15500). K5+/K19-were a gift from Prof. Vilma Band (University of Nebraska). K5+/K19-is a human mammary epithelial cells (HMECs) that has been immortalised with human telomerase reverse transcriptase (TERT)^20^ and were maintained in DFCI media^24^. BRE80-T5 was a gift from Prof. Roger Reddel (University of Sydney). BRE80-T5 is an immortalised mammary epithelial cell line^33^ was maintained in the complete RPMI-1640 medium (Thermo-Fisher Scientific #11875093) containing 10% FBS, 1% L-glutamine, 1% Pen-Strep. BT549, and EFM19 cells were maintained in the complete RPMI-1640 medium (Thermo-Fisher Scientific #11875093) containing 10% FBS, 1% L-glutamine, 1% Pen-Strep. MCF7 and T47D cells were cultured in the complete RPMI-1640 medium (Thermo-Fisher Scientific #11875093) supplemented with 0.01 mg/ml insulin and 2.5% HEPES (Thermo-Fisher Scientific #15630080). CAL120 cell lines were cultures in the complete DMEM medium. All cell lines were kept at 37°C in an incubator containing 5% CO_2_.

### Quantification of global DNA methylation using EPIC array

The Infinium Human Methylation EPIC BeadChip (Illumina, USA) array was performed according to manufacturer’s instructions. To quantify the genome-wide methylation status of bisulfite-converted DNAs. The raw intensity signals obtained from scanning of BeadChips in EPIC were processed with R using the *minfi* package^34^. Methylation CpG (5’-C-phosphate-G-3’) islands were annotated against the genome using the *IlluminaHumanMethylationEPICanno.ilm10b2.hg19* package. Quality of the methylated positions were assessed using the R function *detectionP* from *minfi*^34^ and low-quality positions were discarded. Low quality positions were detected when the methylated and unmethylated channel contained background signal levels (*p-value >0.01*). Data normalization was performed using the R function *preprocessQuantile* from *minfi* and beta-values were generated with the R function *getBeta* from *minfi.* We calculated the fold-change in methylation for each CpG site by comparing the beta-values per CpG between SAM-DNMT3A and cells with SAM-DNMT3A-inactive. Manhattan plots were generated using ggplot2 for each comparison.

### Plasmid constructions

The following plasmids were obtained from Addgene; Plenti-EF1a-SPdCas9-DNMT3B-2A-Blast (dCas9-DNMT3B) (Addgene#71217), pCCL-PGK-SPdCas9-BFP-DNMT1(dCas9-DNMT1) (Addgene#66818) pHRdSV40-NLS-dCas9-24×GCN4-v4-NLS-P2A-BFP-dWPRE (Addgene#60910), pEF1a-NLS-scFvGCN4-DNMT3a (Addgene#100941), pXPR502 (Addgene # 96923), pLenti-EF1a-SPdCas9-DNMT3A-2A-Blast (dCas9-DNMT3A; Addgene#71216), pLenti-EF1a-SPdCas9-DNMT3A(E752A)-2A-Blast (dCas9-inactiveDNMT3A; Addgene# 71218) and pLentiGuide-Puro (Addgene#52963).

For generation of the SAM-DNMT3A sgRNA vector. The transcriptional activator of p65-HSF1 was removed by *KpnI* digestion from pXPR502. The DNMT3A sequence was PCR amplified (PCR primers in Supplementary Table S3) from dCas9-DNMT3A and ligated using gibson ligation. To enable sgRNA cloning via the *ESP3I* restriction site we removed the CGTCTC from the DNMT3A sequence. Using PCR based mutagenesis we generated a point mutation CGTGTC. This point mutation (c.798C>G) in DNMT3A removes the *ESP3I* restriction site without altering the amino acids sequence. To generate the negative control SAM-DNMT3A-inactive sgRNA vector, a gBlock sequence of containing the catalytically inactive DNMT3A inactive was inserted to the *BstXI* and *PstI* sites of SAM-DNMT3A vector.

For cloning of sgRNAs into the sgRNA vectors. For SunTag system sgRNAs were cloned to the *ESP3I* sites of pLentiGuide-Puro (Addgene#52963). For SAM-DNMT3A and SAM-DNMT3A-inactive sgRNAs were amplified by PCR and cloned using golden gate cloning to the ESP3I sites of SAM-DNMT3A or SAM-DNMT3A-inactive vector. All plasmids generated in this study are available through Addgene.

### Cell transfections

HEK293FT cells were transfected with indicated vectors using Lipofectamine3000 (Thermo-Fisher#L3000008). Briefly, 0.5e6 HEK293FT cells were seeded in a six-well plate. 24h post seeding, 1,250ng of each construct were transfected using a 1:1 ratio of DNA to Lipofectamine reagents.

### Quantification of DNA Methylation using pyrosequencing

Genomic DNA (500ng) was bisulfite converted using EZ-DNA Methylation-Gold kit (Zymo-Research#D5006) as per manufactures instructions. The PyroMark Assay Design Software 2 was used to design forward, reverse, and sequencing primers (Supplementary Table S3). PyroMark PCR Kit (Qiagen#978703) was used to amplify genomic fragments surrounding the sgRNA target regions from bisulfite converted DNA according to the company’s instructions. The PCR amplicons were sequenced using PyroMark Q48 Advanced CpG Reagents (Qiagen#974022) based on manufactures instructions.

### Generation of Lentiviral particles

Virus particles were made as previously described^24^. HEK293FT cells (6.5e6) were plated in 10cm dish. After 24h cells were transfected with 8,750ng of indicated vector and 8,750ng of psPAX2 (Addgene#12260) and 875ng of pMD2.G (Addgene#12259) in Opti-MEM (Thermo-Fisher#51985091) (final volume 500μl) and 103µl of Lipofectamine 3000. Supernatant was collected at 48h and 72h post transfection and 30% of FBS was added. Virus particles were kept at –80°C.

### Transduction of cells with SAM-DNMT’s

To induce DNA methylation using *SAM* system cells were infected with a lentiviral vector expressing dCas9-DNMT3A and a modified sgRNA lentiviral vector (Fig. 1A). 1.2e6 cells were plated in 8 ml media and supplemented with 0.5μg/ml of polybrene in a 10cm dish and 2ml of dCas9-DNMT3A expressing virus was added to the media. 24h later, media was removed and replaced with a media containing 10μg/ml of blasticidin (Thermo-Fisher#A1113903). Following selection (7 days), cells were maintained and expanded in a media containing 5μg/ml of blasticidin for later application.

To express an sgRNA in these cells, 3e5 cells/well of dCas9-DNMT3A expressing cells were plated on a 6-well plate. 24h later media was replaced with 1.5ml of media containing 0.5μg/ml of polybrene and 500μl of modified sgRNA lentiviral vector. Following 24h, the media was replaced with media containing 5μg/ml of blasticidin and 2μg/ml of puromycin. After selection (2 days) puromycin was reduced to 1μg/ml.

### Generation of pooled sgRNA libraries

Pooled sgRNA libraries were generated as previously described^35^. For each of heritable methylation marks, 10 sgRNAs were designed (Supplementary Table S1). Each sgRNA had *ESP3I* cut sites and flanking PCR handles (AGGCACTTGCTCGTACGACGCGTCTCACACCG[20ntsgRNA]GTTTCGAGACGTT AAGGTGCCGGGCCCACAT-3’). PCR amplification was performed using primers (Fwd: 5’-AGGCACTTGCTCGTACGACG-3’, Rev: 5’-ATGTGGGCCCGGCACCTTAA-3’). Golden gate cloning was applied to insert the library into the *ESP3I* sites of SAM-DNMT3A or SAM-DNMT3A-inactive. Library representation was verified by NGS sequencing.

### SAM-DNMT3A screen

MCF7, BRE80-T5 and T47D cells stably expressing dCas9-DNMT3A or dCas9-DNMT3A-inactive were transduced with the sgRNA pooled library virus at an MOI of 0.3. 24h later infected cells were selected using Puromycin (2μg/ml). Cells were maintained with blasticidin and puromycin throughout the screen, to ensure expressions. 21 days post infection, DNAs were extracted using NucleoSpin Blood XL (Macherey-Nagel#740950). Samples were prepared for sequencing as previously described^35^.

### Crystal violet proliferation assay

2e4 cells/well were plated on a 24-welll plate in triplicates and allowed to propagate for 7-10 days. Cells were washed with Dulbecco’s phosphate-buffered saline (DPBS; ThermoFisher#14190-144) and 1ml of 10% formalin was added into each well and incubated for 20min at room temperature. After removal of formalin, 0.5ml of 0.5% (w/v) crystal violet (Sigma#C0775-25G) was added. After 20min, crystal violet was removed, and plates were washed with water. For quantification, 0.5ml of 10% Acetic acid was added to each well and incubated for 30min. 100µl of the dissolved solution was added to a 96-well plate and quantified by measuring the OD at 570nm.

### Analysis of global DNA methylation at LINE-1 using the HRM assay

7 days post infection with the indicated sgRNAs, genomic DNA was extracted using DNeasy Blood and Tissue Kit (Qiagen#69504) and bisulfite converted using EZ-DNA Methylation-Gold kit. CpGenome™ Human Methylated & Non-Methylated DNA standard set (Sigma#S8001M) was used to prepare a set of methylation standards (0%, 25%, 50% and 100% methylated DNA standards). The PCR amplification of all bisulfite converted DNA was performed in a 20μl reaction volume containing 1μl of bisulfite converted DNA, 10μl of MeltDoctor HRM Mastermix (ThermoFisher#4415440), 1μl of forward and 1μl of reverse primers (10μM) and 7μl of H_2_O. We targeted LINE-1 region to measure global DNA methylation using primer F: 5’-GCGAGGTATTGTTTTATTTGGGA-3’; R: 5’-CGCCGTTTCTTAAACC-3’ (Tse et al (2011)). The HRM analysis was initiated by one cycle at 95°C for 10min followed by 40 cycles at 95°C for 15sec, 60°C for 1min. The melting process was followed by denaturing step at 95°C for 10sec, annealing at 60°C for 1min and by a stepwise increase in temperature from 60°C to 95°C, with a rate of 0.1°C per second. Finally, The HRM results were analysed using the Thermo-Fisher HRM software.

## Acknowledgments

This work was supported by an NHMRC Synergy grant to J.R. and M.C.S (grant number: 2011329). J.R. is supported by a Victoria Cancer Agency fellowship (grant number: MCRF20035). We thank the Functional Genomics Platform, and the Genomics and Bioinformatics Platform at Monash University for help with loss of function screens and data analysis. We thank Dr. Gavin Knott (Monash University) for helpful discussions. We thank Prof. Georgia Chenevix-Trench (QIMR), Prof. Roger Reddel (University of Sydney) and Prof. Vilma Band (University of Nebraska) for providing cell lines.

## Supplementary Figures

**Supplementary Figure 1:**
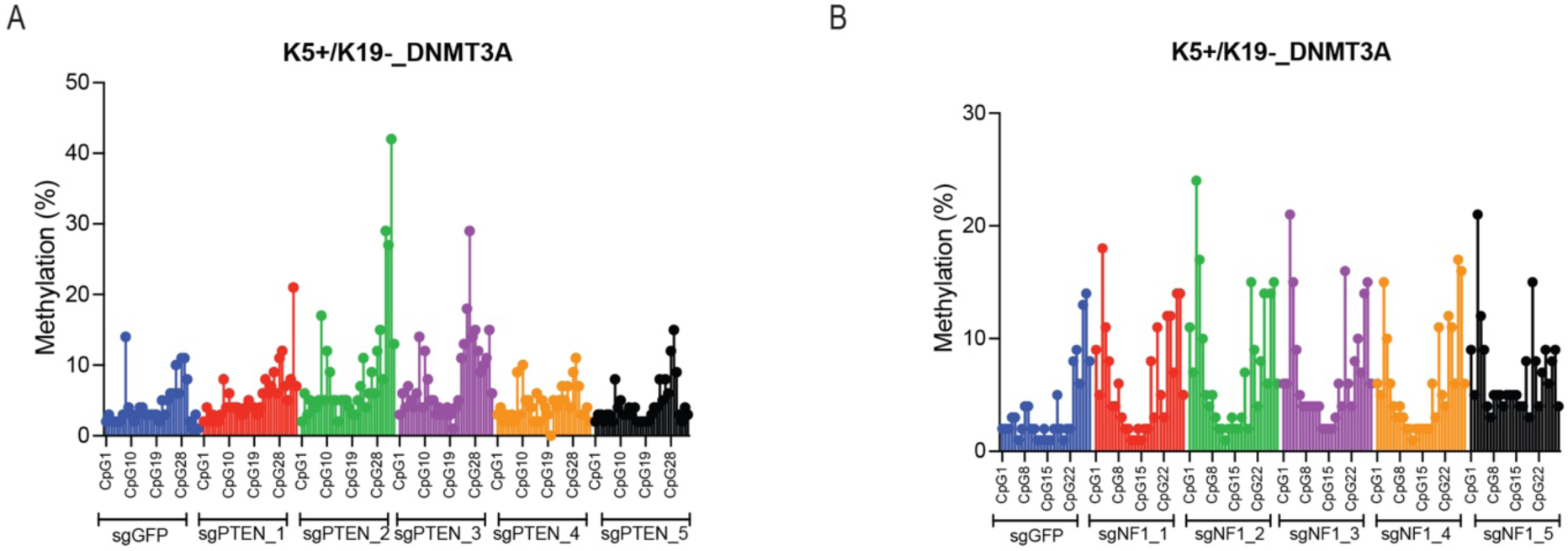
SAM-DNMT3A induces high levels of DNA methylation at desired sites. Pyrosequencing was used to measure DNA methylation at CpG sites in of K5+/K19-^20^ cells 7 post transduction with SAM-DNMT3A system. (A) For the *PTEN* promoter. (B) For *NF1* promoter.

**Supplementary Figure 2:**
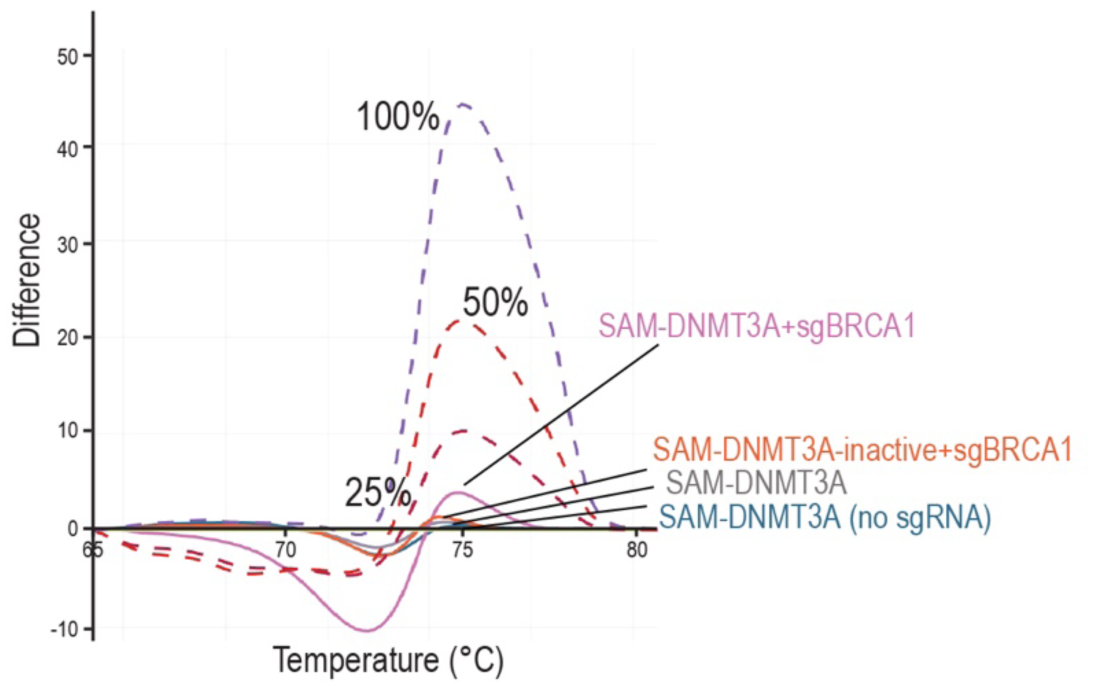
SAM-DNMT3A pooled screens do not identify any hits associated with proliferation. HRM assay using *BRCA1* probes in HEK293T cells expressing SAM-DNMT3A or SAM-DNMT3A-inactive with a *BRCA1* targeting sgRNA.

**Supplementary Figure 3:**
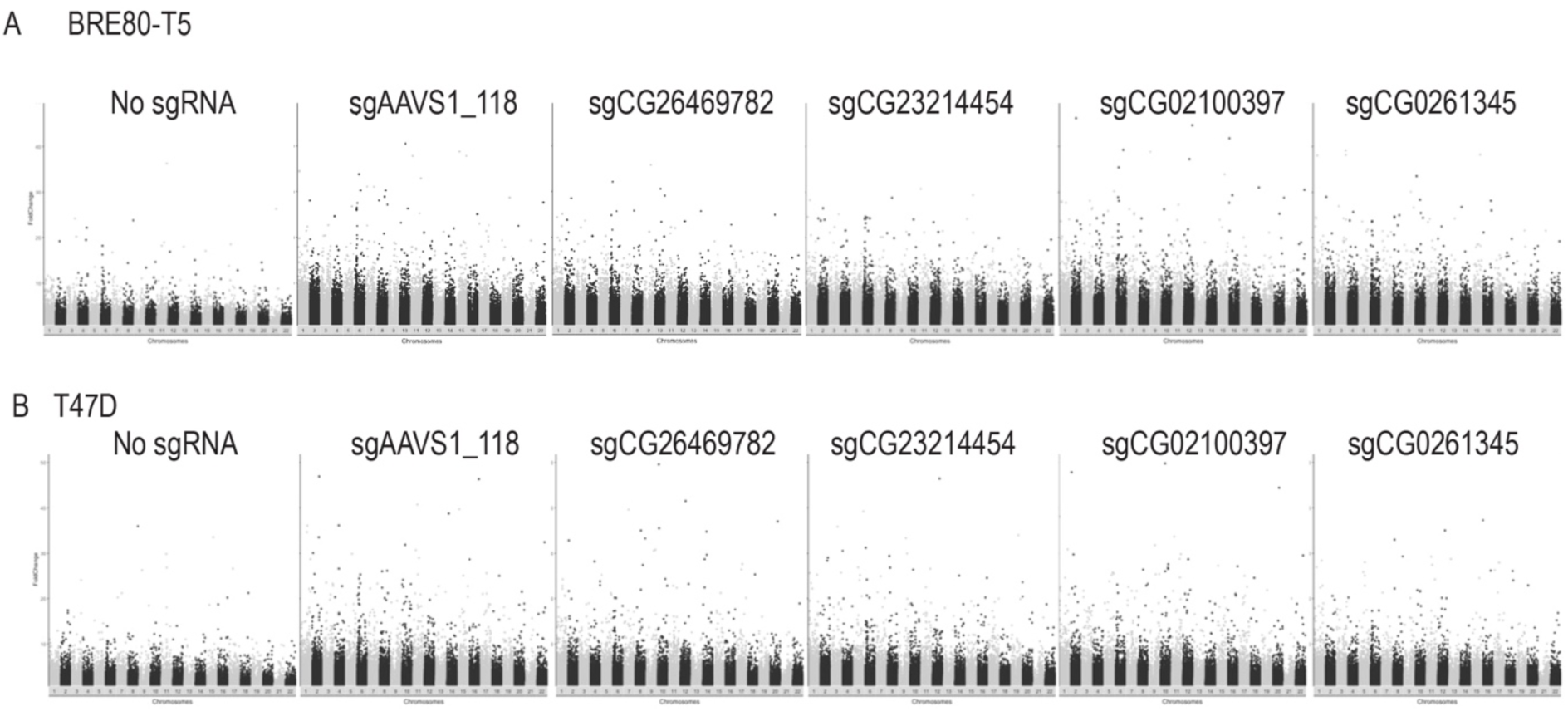
SAM-DNMT3A induces global non-specific DNA methylation. (A) Manhattan plot showing fold changes in DNA methylation obtained by comparing DNA methylation levels (from EPIC arrays) in cells expressing the same sgRNA with SAM-DNMT3A and SAM-DNMT3A-inactive from with different sgRNA treatments in BRE80-T5. (B) Like (A) in T47D cells.

**Supplementary Figure 4:**
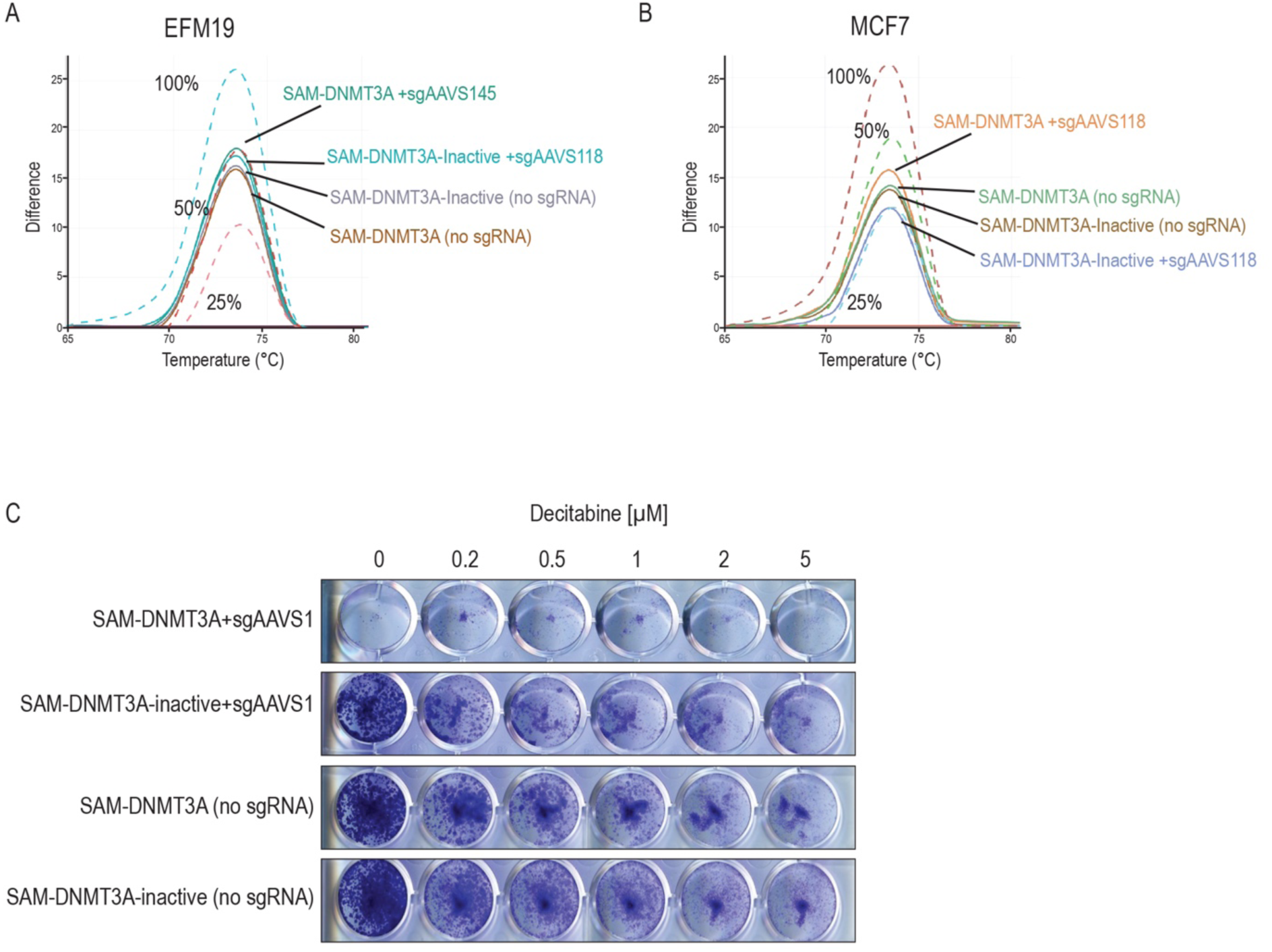
Induction of DNA methylation is a vulnerability in ER positive breast cancers. HRM assay quantifying DNA methylation at LINE1 sites. Dotted lines are standards containing the indicated percent of methylated DNA made from a mix of methylation and unmethylated DNA. (A) EFM19 (B) MCF7. (C) Crystal violet images of MCF7 cells expressing SAM-DNMT3A or SAM-DNMT3A-inactive with or without an AAVS1 targeting sgRNA treated with increasing concentrations of the DNMT inhibitor decitabine.

## Supplementary Tables Legends

**Supplementary Table S1: sgRNAs used in SAM-DNMT3A screens.**

**Supplementary Table S2: Raw reads sgRNA and gene fold changes in pooled SAM-DNMT3A screens.**

**Supplementary Table S3: Primers and sgRNA sequences used in this study.**

